# Developmental genetics in a complex adaptive structure, the weevil rostrum

**DOI:** 10.1101/287185

**Authors:** S.R. Davis

## Abstract

The rostrum of weevils (Curculionidae) is a novel, complex, adaptive structure that has enabled this huge beetle radiation to feed on and oviposit in a wide spectrum of plant hosts, correlated with diverse life histories and tremendous disparity in rostrum forms. In order to understand the development and evolution of this structure, transcriptomes were produced in *de novo* assemblies from the developing pre-pupal head tissues of two distantly related curculionids, the rice weevil (*Sitophilus oryzae*) and the mountain pine beetle (*Dendroctonus ponderosae*), which have highly divergent rostra. While there are challenges in assessing differences among transcriptomes and in relative gene expression from divergent taxa, tests for differential expression patterns of transcripts yielded lists of candidate genes to examine in future work. RNA interference was performed with *S. oryzae* for functional insight into the Hox gene *Sex combs reduced. Scr* has a conserved function in labial and prothoracic identities, but it also demonstrates a novel role in reduction of ventral head structures, namely the gula, submentum, and associated sulci, in weevils. Ultimately, this study makes strides towards elucidating how the weevil rostrum initially formed and the profound phenotypic diversity it has acquired throughout the curculionoid lineages. It furthermore initiates a better understanding of the genetic framework that permitted the diversification of such an immense lineage as the weevils.

**Summary statement:** This study begins exploring the development of a novel, complex structure in one of the largest families of organisms, the weevils.

## Introduction

Weevils (Coleoptera: Curculionoidea) are a diverse group of extant organisms with nearly 60,000 described species (Anderson, 1995). They are of great agricultural significance because they are associated with all major groups of plants and plant tissues. The weevil rostrum, for example, is a key evolutionary innovation that has enabled this group to feed on and oviposit in nearly all plant tissues, giving rise to diverse life histories and tremendous diversity in rostrum form (Fig. 1). Insights into comparative development of the rostrum will provide a better understanding of its evolutionary origins and modification, and may help to illuminate the diversification of the lineage (Anderson, 1995, Oberprieler et al., 2007). In beetles (Coleoptera), *Tribolium* (Tenebrionidae) has been the primary model in which to study evolutionary developmental mechanisms (Tomoyasu et al., 2005). Exciting and relatively recent additions include a species of *Onthophagus* (Scarabaeidae), which has emerged as a new model for examination of horn development (Moczek and Nagy, 2005; Moczek and Rose, 2009; Moczek et al., 2006, 2007), as well as other beetle species for similar examinations in exaggerated male traits (e.g., Ozawa et al., 2016). Although weevils are an enormous group and many species are significant agricultural pests, no weevil species have been utilized in developmental studies concerning phenotypic evolution; however, somewhat related studies have examined weevil olfactory genes (Bin et al., 2017), RNAi efficacy in weevils (Christiaens et al., 2016; Prentice et al., 2016), as well as microbiome contributions in weevil development (e.g., Anbutsu et al., 2017; Heddi et al., 1999; Kuriwada et al., 2010; Vigneron et al., 2014).

**Fig. 1.**
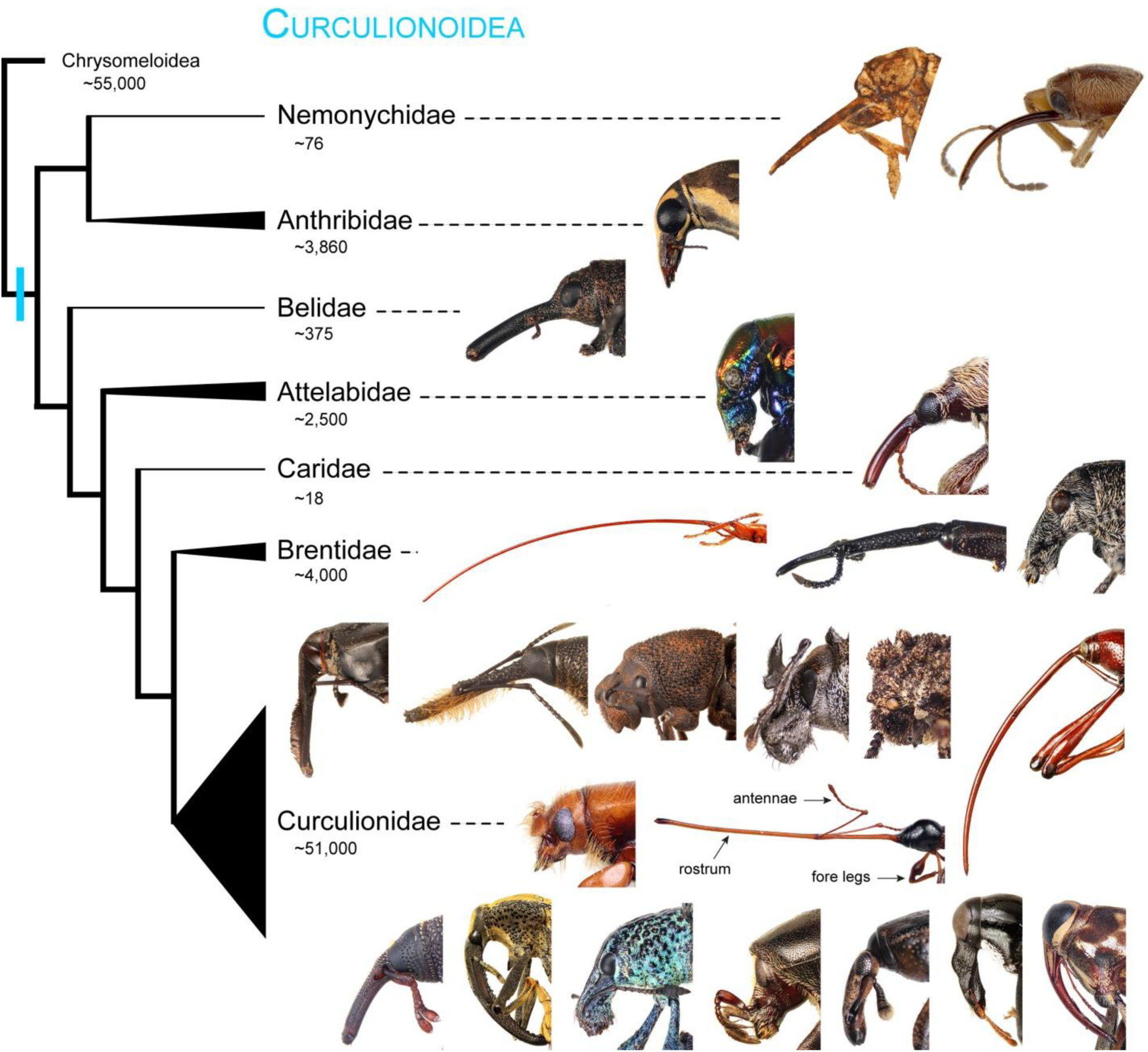
Current phylogenetic summary of the weevil superfamily Curculionoidea, depicting heads of various taxa within each family displaying a diversity of rostral lengths and forms. Approximate species diversity (from Oberprieler *et al.* 2007) within each lineage is listed below the clade names. All weevil heads depicted are of extant taxa excepting the first left image in Nemonychidae of a Late Jurassic Karatau compression fossil. Relationships based on Davis (2017), Mckenna *et al.* (2009), and Oberprieler *et al.* (2007).

From previous morphological and histological observations (Davis, 2011), patterning of the adult rostrum begins in the early 4^th^ (last) instar larva and continues throughout the pupal stage. Aside from adult appendage patterning, relatively little research has focused on the formation of adult structures in insects. Much of this work has been done relatively recently (see Jockusch and Smith, 2015 for general review), examining diverse features such as horn development (as mentioned above), mouthparts (e.g., Angelini et al., 2012), metamorphosis (e.g., Konopova and Jindra, 2008), head and body wall patterning (e.g., Smith and Jockusch, 2014), and wings (e.g., Lozano and Belles, 2014; Martin and Reed, 2010; Ohde et al., 2013; Prud’homme et al., 2011; Shimmi et al., 2014; Tomoyasu et al., 2005, 2009). Because head segmentation is established in the embryo, the majority of research has traditionally focused on these early stages to understand developmental pathways. The weevil head, however, has diverged from the typical beetle head plan to accommodate a pre-ocular extension of the anterior portion of the head capsule, producing a rostrum with the mouthpart appendages at the apex. The rostrum incorporates all head segments and largely the sclerites of the parietals, pleurostoma, gena, occiput, postgena, postmentum, and frons. These head regions are demarcated mostly by cranial lines, sulci, tentorial pits, and landmark features such as the eyes, antennae, and mouthparts (Fig. 7). These segments can be delimited posteriorly with the homeodomain segment polarity gene *engrailed* (*en*), which is known to specify posterior segment boundaries in *Drosophila* and *Tribolium* (Rogers and Kaufman, 1996; Schmidt-Ott and Technau, 1992; Schmidt-Ott et al., 1994). Also, since the rostrum is formed from the elongation of different combinations of these segments, its development may combine genetic elements of appendage patterning and segment identity. In terms of the latter, a reasonable start is to examine members of the Antennapedia Hox gene complex. This study addresses the function of *Sex combs reduced* (*Scr*) in weevil rostrum development.

Additionally, transcriptomes were produced in *de novo* assemblies from the developing head tissues of two weevil species, which are major pests, *Sitophilus oryzae* (the rice weevil) and *Dendroctonus ponderosae* (the mountain pine beetle), the latter of which has had a genome produced (Keeling et al., 2013). Both species are considerable pests in stored grains and conifer forests, respectively, representing disparate clades and divergent rostral forms. More specifically, *Sitophilus* possesses a rostrum of moderate length, while *Dendroctonus* essentially has no rostrum, in which it presumably was lost based on its derived phylogenetic position within Curculionidae. While there are difficulties in assessing differences among transcriptomes from divergent taxa, tests for differential expression patterns of transcripts were performed to identify genes involved in the initial acquisition of a rostrum and subsequent diversification. Ultimately, the goal of this and following studies is to characterize the genetic differences that produced the profound phenotypic diversity observed in the rostrum and to better understand the genetic framework that permitted the diversification of such an immense lineage as the weevils. To begin, functional tests of candidate genes were performed through larval RNA interference (RNAi). Weevil head regions and anatomical terminology follow that in Davis (2011, 2017). The hypothesized adult head regions of wild-type *S. oryzae* have been slightly modified in this study from their original presentation in Davis (2011) following more in-depth study of the adult anatomy.

## Materials and Methods

### Animal husbandry

The *Sitophilus oryzae* colony is maintained on whole barley in circular plastic containers, ∼7cm in height and ∼10 cm in diameter, with small air holes punctured in the lids. As the immature stages are endophytic, larvae develop and pupate within the barley kernels and desired stages are carefully extracted using a razor blade. *Dendroctonus* larvae were collected from felled logs.

### Electron microscopy

Scanning electron microscopy (SEM) images were captured using a LEO 1550 FESEM (Field Emission Scanning Electron Microscope). Specimens and dissections were mounted on aluminum SEM stubs using carbon conductive tape and coated with gold.

### Histological sectioning

Both paraformaldehyde-fixed and ethanol-fixed heads were used. After tissues were dehydrated in 100% ethanol, infiltration of head tissue consisted of approximately 2-hour incubation periods through a series of 1:1 then 1:2 ethanol to LR White® (an acrylic resin) mixtures. Heads were placed in gelatin capsules and embedded in pure LR White®. Following thermal curing in an oven for 24 hours at 60° C, embedded heads were removed from the capsules and sectioned using a Leica EM UC6 ultratome and diamond knife, producing semithin sections ∼5–6 μm thick. Sections were transferred to glass slides, stained in toluidine blue, air dried, mounted in Permount™, and digitally photographed. A z-stack was acquired of several photomicrographs using the software CombineZ, and edited in Adobe Photoshop CS3.

### Transcriptome assembly, annotation, and DE analysis

Pre-pupae (pharate pupae) were obtained for two curculionid species: *Dendroctonus ponderosae* and *Sitophilus oryzae* (Fig. 2). Ten frozen *S. oryzae* pre-pupae and seven *D. ponderosae* pre-pupae were decapitated and the pooled RNA extracted from the heads using Trizol. Library preparation followed the Illumina TruSeq protocol, incorporating Covaris shearing as an alternative to chemical shearing and excluding the cDNA DSN Normalization step, and Illumina 100bp, paired-end sequencing was performed on a HiSeq2000 platform for one library of each species. Separate transcriptome assemblies were performed *de novo* using Trinity. As two fairly distantly related species were being compared, reciprocal best BLAST hits were determined using the conditional reciprocal best BLAST (CRB-BLAST) implemented by Aubry et al. (2014) to assign orthologs. Using the reduced transcriptomes, which retained those transcripts identified as orthologs using CRB-BLAST, Bowtie v2 (Langmead and Salzberg, 2012) and RSEM v1.1.17 (Li and Dewey, 2011) were used for read mapping and transcript quantification; Salmon (Patro et al., 2017) was also implemented as a comparison. Sequence reads were mapped and quantified onto their respective reduced transcriptomes, then the raw counts were used as input into edgeR (Robinson et al., 2010) for differential expression (DE) analysis. As no RNAseq replicates were obtained, DE analysis in edgeR was performed through manual adjustment of the Biological Coefficient of Variation (BCV) set at 0.2, 0.4, and 0.6. As recommended when using DE software without replicates, no statistical analyses were performed. Instead, differential expression levels were explored and compared through iterations of modifying the BCV.

Blast2GO v3 (Conesa and Götz, 2008) and NCBI Blast 2.2.25+ were used to obtain DE transcript identities and were blasted to the *Drosophila melanogaster* CDS database Dmel_r6.18. Transcripts were also compared to the UniProt database for obtaining biological class Gene Ontology (GO) ID’s and to the PANTHER classification system (Mi et al., 2013) for obtaining protein class annotations.

**Fig. 2.**
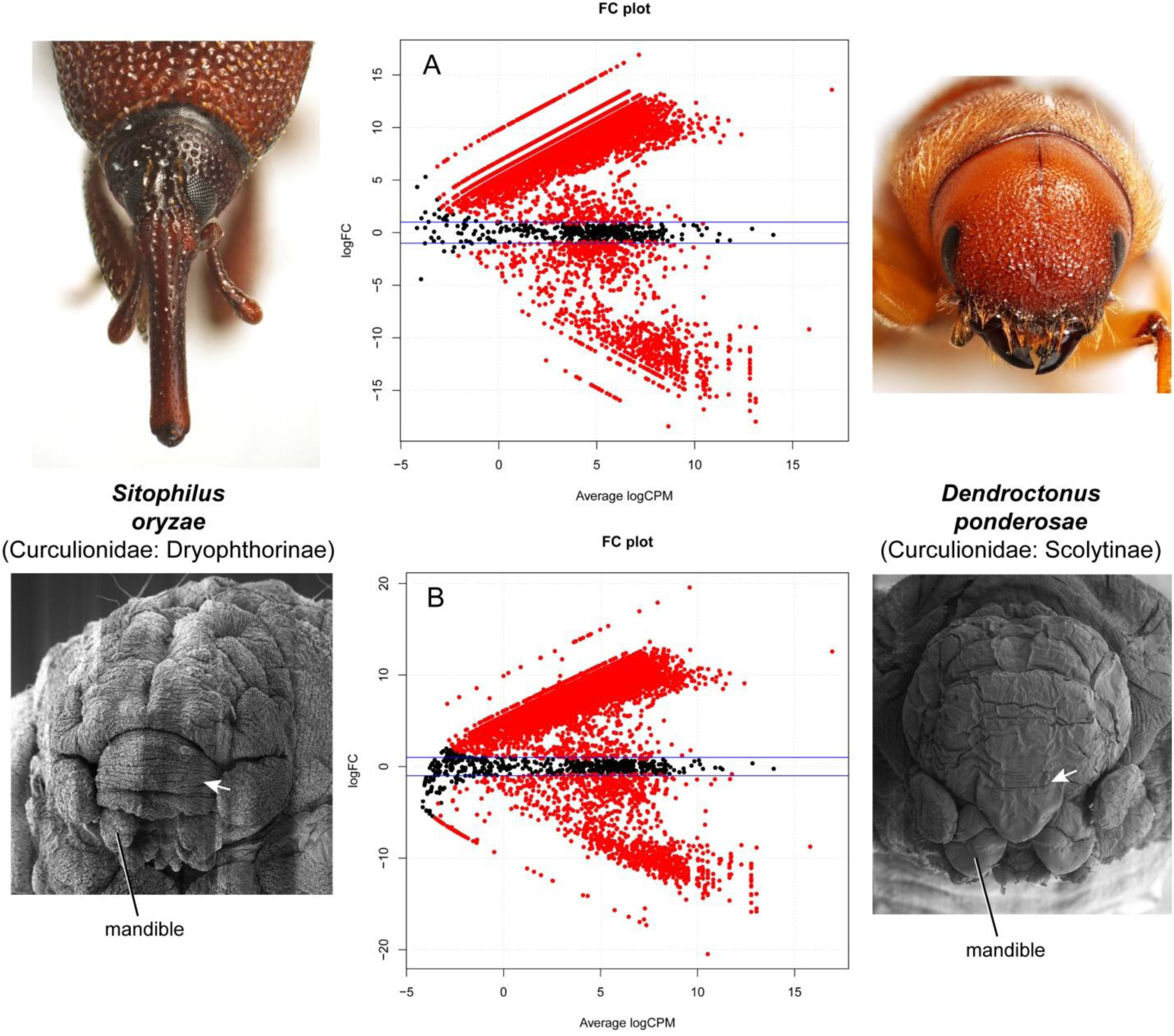
MA (Fold Change) plots generated in edgeR with BCV=0.2. **A**, DE analysis using the RSEM transcript quantification method; **B**, DE analysis using the Salmon transcript quantification method. Log-Fold Change on the y-axis and log-Concentration (in Counts Per Million) on the x-axis, depicting DE transcripts in red. Blue threshold lines indicating fold-change of 2. The comparison was made between transcriptomes and RNA-seq data generated from the pre-pupal (pharate pupal) heads of *S. oryzae* (left side) and *D. ponderosae* (right side), of which the pre-pupal (SEM) and adult heads are shown; arrows on SEM’s point to epithelial folds in rostrum or lack thereof.

### RNA interference

RNA interference (RNAi) was performed to create loss-of-function phenotypes generally following the protocols for *Tribolium* of Tomoyasu and Denell (2004) and Tomoyasu et al. (2005, 2009). Briefly summarized, an *Scr* gene fragment of 427 bp was amplified using the primer pair 5’-TTACTCGCCTCCCCAAGTAG-3’ and 3’-CGGCGAGGACAAATTCTAAG-5’ and cloned into the pCR™4-TOPO® plasmid using the TOPO® TA cloning® kit (Invitrogen). Double-stranded RNA (dsRNA) was produced *in vitro* using the Ambion MEGAscript kit following the manufacturer’s protocol. Using pulled glass capillary needles, 1 μL dsRNA (4 μg/μL) was mixed with green food coloring and injected into 4th instar larval *Sitophilus* specimens (larval RNAi) using ice packs to immobilize the specimens. Negative control injections consisted of water mixed with green food coloring and *Scr* dsRNA produced for *Tribolium castaneum*.

## Results and Discussion

### Transcriptome assembly, Read mapping, and DE analyses

In determining which tissues to sample for transcriptome acquisition, whole heads were used (as opposed to discrete rostrum tissue) on account of the extreme rostrum reduction in *D. ponderosae*. As is visible in SEM’s of late pre-pupal heads of the two taxa (Fig. 2), many epithelial folds are present in the developing rostrum of *S. oryzae* while there are none in that region of *D. ponderosae*. Transcriptome assembly statistics from Trinity are provided in Table 1. Just over three times as many transcripts were identified in *D. ponderosae* than in *S. oryzae*. After read mapping, these superfluous contigs were identified mostly as incorrectly assembled transcript isoforms. To account for excessive isoform generation, some assembly statistics, in addition to being calculated using all of the contigs, were also calculated in Trinity using the “longest isoform per Trinity gene,” resulting in more comparable assembly numbers.

**Table 1.** Trinity de novo transcriptome assembly statistics for *Sitophilus oryzae* and *Dendroctonus ponderosae*.

|  | *Sitophilus oryzae* |  | *Dendroctonus ponderosae* |  |
| --- | --- | --- | --- | --- |
|  | all contigs | longest isoform per 'gene' | all contigs | longest isoform per 'gene' |
| Total Trinity 'genes' | 34,843 |  | 86,308 |  |
| Total Trinity transcripts | 59,972 |  | 172,316 |  |
| GC content | 35.40% |  | 38.65% |  |
| Contig N20 (bp) | 3,805 | 3,427 | 12,591 | 4,161 |
| Contig N50 (bp) | 1,744 | 1,562 | 8,035 | 1,050 |
| Average contig length (bp) | 902 | 810 | 2,195 | 643 |

Following assembly, reciprocal best BLAST hits were obtained to identify orthologous transcripts from which to conduct further analyses, a process that resulted in a reduction of comparable transcripts. Specifically, the number of transcripts in the reduced dataset was 8,912 for both taxa. This reduction at first seems sizeable in comparison to the total number of Trinity contigs for each taxon. When comparing these totals to the proportions of the contigs which obtained BLAST hits to the genomes of both *D. melanogaster* and *T. castaneum*, only 32% of both transcriptomes had gene annotations. Therefore, it is assumed that at least half of the contigs were either incomplete or improperly assembled. While it is assumed that information is lost in dataset reduction, thereby influencing downstream differential expression (DE) analyses, it likely is rare that all transcripts representing a certain gene are eliminated. The Illumina reads were then mapped onto the respective reduced transcriptomes containing one to one orthologs and transcript counts were quantified. Two different methods were used to perform this step in order to observe any differences in DE analysis outcome: RSEM, an alignment based method which utilizes Bowtie2 for mapping, and Salmon, which was set for quasi-mapping. The count data was used as input for DE analysis in edgeR, and three BCV values were used to assess transcript-fold differences with increasing dispersion values (from low to moderate) since no biological replicates were obtained (Fig. 2). DE results using RSEM and Salmon were relatively comparable, with the total number of down-regulated transcripts being six to seven times greater than the total of those up-regulated, and the total number of DE transcripts slightly increased using the former method (Table 2).

**Table 2.** Results obtained using two transcript quantification methods, RSEM and Salmon, and using various BCV values for differential expression (DE) analysis with edgeR. Values refer to number of Trinity transcripts identified as being differentially expressed (up- or down-regulated) or not.

|  | **RSEM** |  |  | **Salmon** |  |  |
| --- | --- | --- | --- | --- | --- | --- |
|  | BCV=0.2 | BCV=0.4 | BCV=0.6 | BCV=0.2 | BCV=0.4 | BCV=0.6 |
| down-regulated | 1098 | 931 | 802 | 1363 | 1179 | 1004 |
| not DE | 423 | 750 | 1024 | 557 | 881 | 1231 |
| up-regulated | 7391 | 7231 | 7086 | 6992 | 6852 | 6677 |

After exploring expression levels at different levels of data dispersion, a conservative perusal of DE transcripts was conducted by using those identified from edgeR at BCV=0.2 and using Salmon for contig quantification, which was a total of 8,355 transcripts. These transcripts were blasted to the *D. melanogaster* CDS database and compared to the UniProt and PANTHER databases for obtaining Gene Ontology (GO) ID’s (Fig. 3A) and protein class annotations (Fig. 3B), respectively. Of those transcripts, a list of some interesting transcripts was produced representing many genes which are hypothesized to be involved in rostrum development and differentiation (Table 3) and which will be the topic of subsequent study. Those genes of notable mention include *Sex combs reduced* (*Scr*), *homothorax* (*hth*), several *broad* (*br*) isoforms in the *Broad-Complex* (*BR-C*), *expanded* (*ex*), *split ends* (*spen*), *Hairless* (*H*), and *Dorsal switch protein 1* (*DSP1*).

**Table 3.**
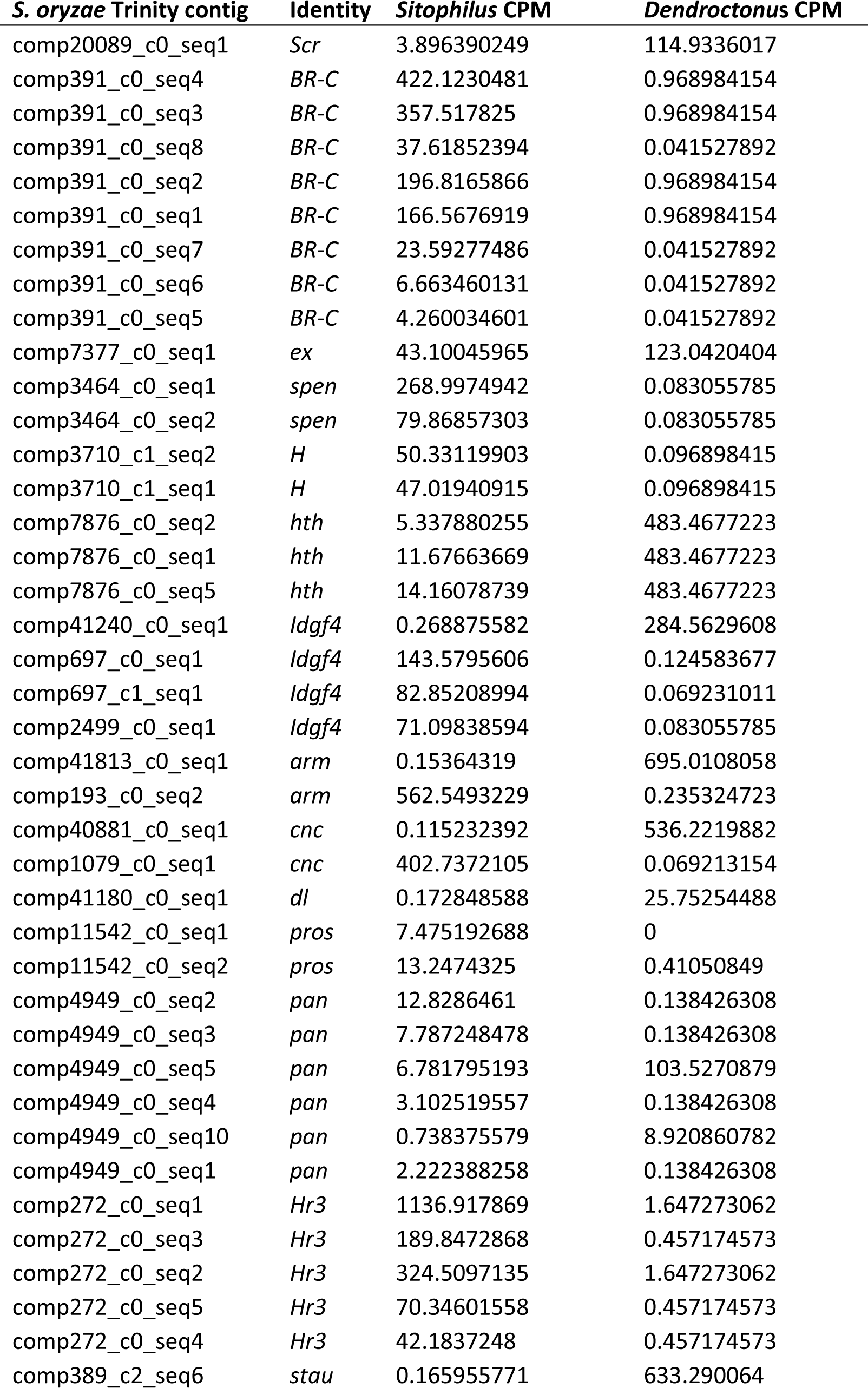
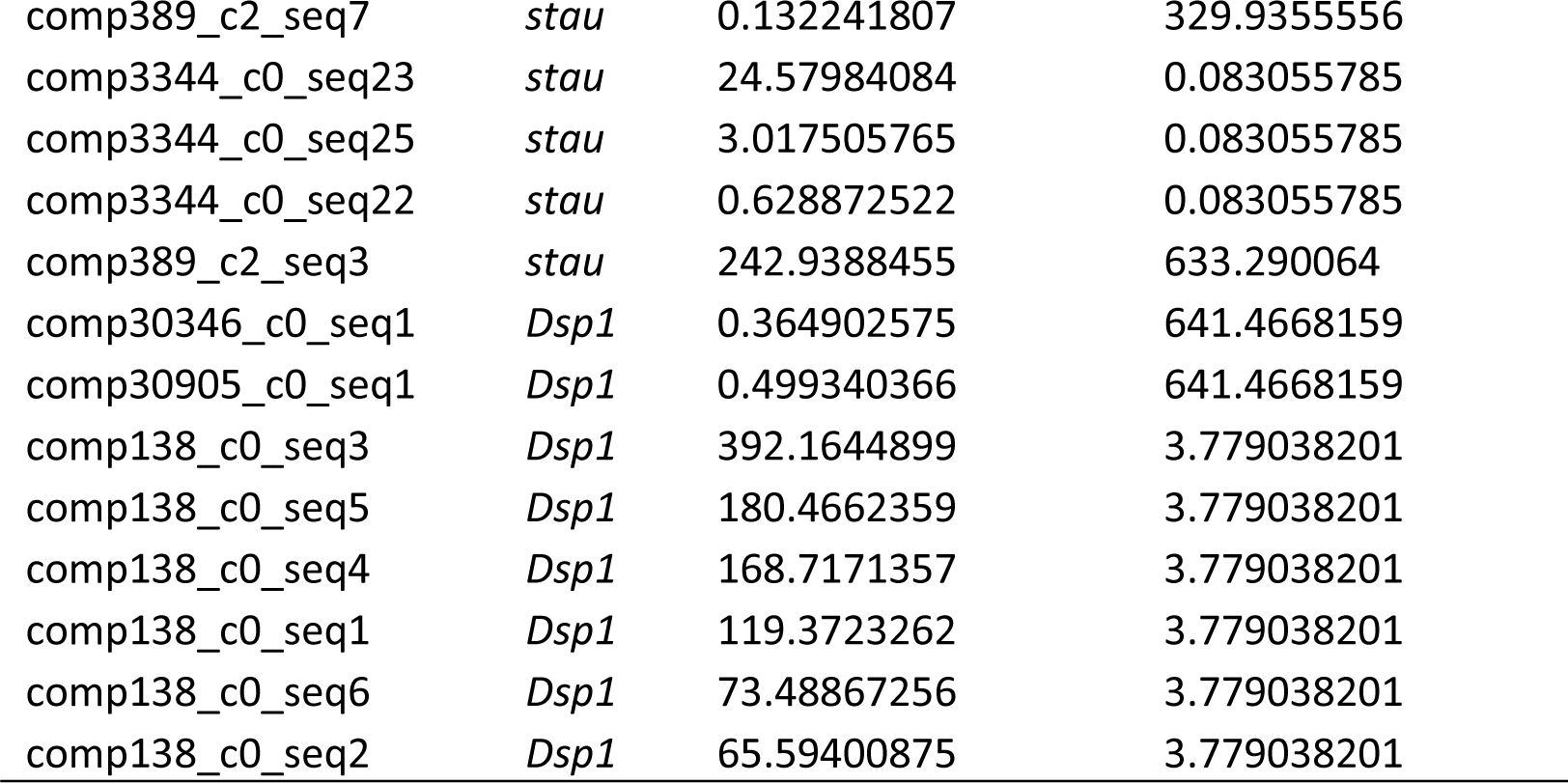
List of interesting transcript isoforms filtered from EdgeR analysis using BCV=0.2. CPM=counts per million.

**Fig. 3.**
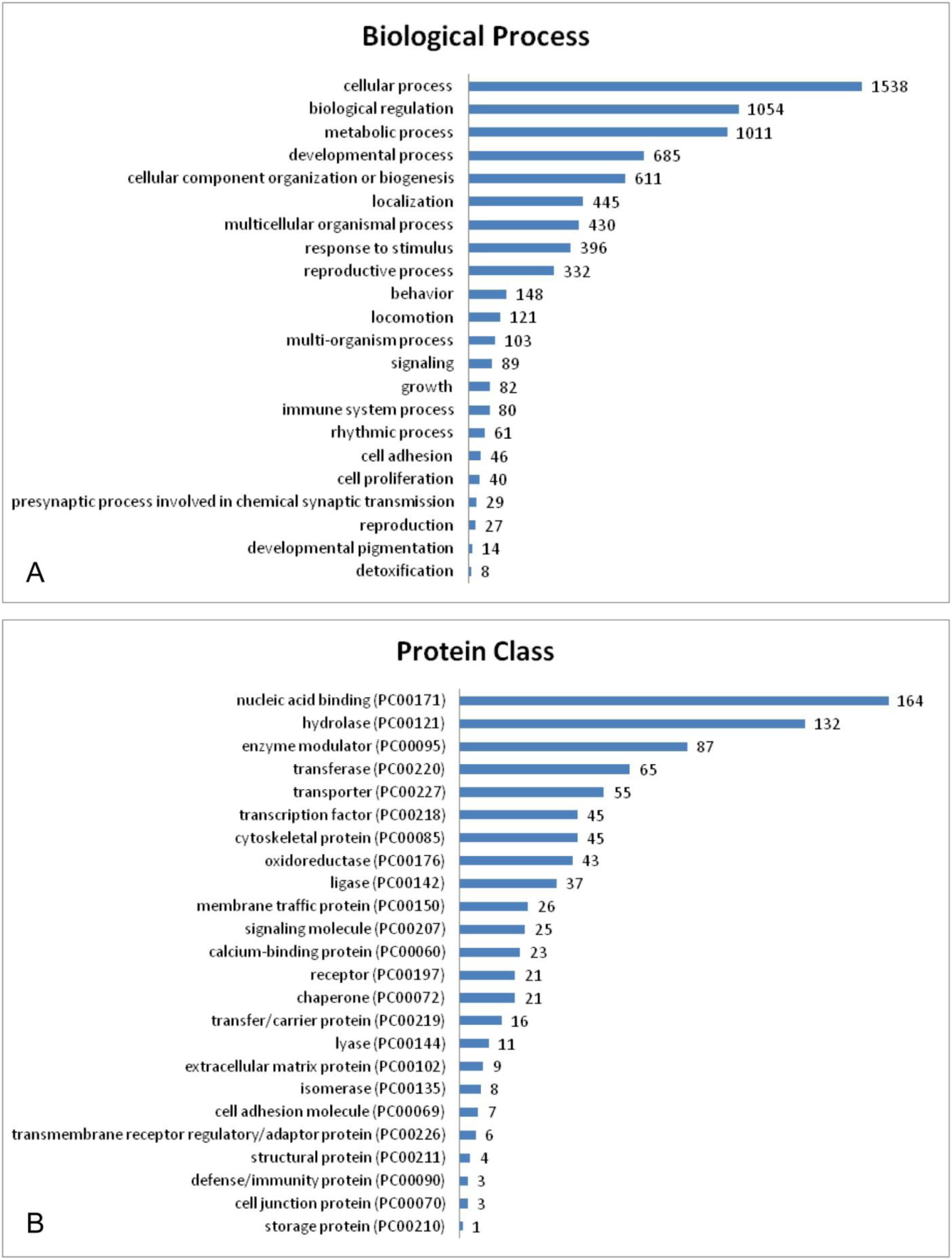
Annotation of DE transcripts obtained from edgeR analysis with BCV=0.2 and using Salmon for transcript quantification. **A**, biological process Gene Ontology (GO) terms obtained from UniProt database; **B**, protein class ID’s obtained from the PANTHER classification system.

From Salmon transcript quantification and edgeR DE analysis, *Scr* expression is higher in *D. ponderosae* than in *S. oryzae* (Table 3). The rostrum in the wild-type of *D. ponderosae* is strongly reduced (qualitatively absent), the prementum slightly shorter, and the postmentum markedly shorter. Other than these differences, the external morphology of the wild-type head is not drastically different from that of *S. oryzae*. In other words, the much lower expression of *Scr* in the latter taxon does not appear to have much influence on the rostral phenotype with exception perhaps to the length of parts of the labium and its posterior regions. In adult insects, *Scr* seems to play a conserved role in the identities of the labium and prothorax, in which mutants display labial to maxillary and prothoracic to mesothoracic homeotic transformations. In *T. castaneum Scr*-RNAi, parts of the labium assume maxillary morphology and the prementum and mentum are reduced in size (Smith and Jockusch 2014).

Similar to *Scr*, the expression levels of *hth* are higher in *D. ponderosae*. The complex interactions of *hth* with *extradenticle* and other Hox proteins can cause segmental identity transformations and appendage patterning malformations. In *T. castaneum hth*-RNAi, there are various malformations in the labium and maxillae, and the prementum and mentum are reduced in size (Smith and Jockusch, 2014).

The expression of several *br* isoforms, all at higher levels in *S. oryzae*, indicates that this complex perhaps has a distinct role in rostrum formation. It acts downstream of juvenile hormone and is an early ecdysone-inducible gene that encodes a family of DNA binding proteins, one of which, *br*, has multiple isoforms that are expressed during metamorphosis and are responsible for tissue-specific morphogenesis (Karim et al., 1993; Konopova and Jindra, 2008; Parthasarathy et al., 2008; Suzuki et al., 2008). Malformations caused by *br*-RNAi can be severe depending on the tissue and it can hinder pupation. Preliminary examination of one isoform in *S. oryzae* has thus far demonstrated a function corresponding to the expression levels found in this study and reduces proliferation of tissue in the rostrum (unpublished results).

*Expanded* is a Hippo tumor suppressor pathway component involved in imaginal disc development and cell proliferation. Its expression is greater in *D. ponderosae*, suggesting its function may involve suppression of rostrum growth. The function of *ex* has not been observed in *T. castaneum*, but in *D. melanogaster* it is necessary for proper imaginal disc growth and mutants can show tissue hyperplasia or degeneration. Therefore, due to its association with disc growth, its function in the weevil rostrum is perhaps to limit its extension in cases of higher levels of expression such as in *D. ponderosae*.

Effects of s*plit ends* are seen in cell-fate specification in neurons and with homeotic segment identity as well. In concert with Hox proteins, it is required for the differentiation and sclerotization of certain larval and adult head regions (Wiellette, 1999). As the *Drosophila* larval head is greatly reduced in comparison to the majority of other holometabolous insects, the role of *spen* in the weevil head and rostrum may be of some importance. Its expression is shown to be much higher in *S. oryzae*, indicating a role perhaps in rostrum sclerite elongation.

*Hairless* is mostly known for its roles in bristle patterning, affecting sensory organ precursor cell fate through Notch pathway inhibition (Bang et al., 1991), as well as in imaginal disc development. When overexpressed, *H* induces apoptosis through down-regulation of EGFR signalling activity (Protzer et al., 2008). In *S. oryzae* it is up-regulated, and perhaps its expression is involved in the eye-antennal disc.

*DSP1* is a corepressor of *dorsal*, a protein known for its involvement in dorso-ventral polarity in developing *D. melanogaster*. Loss of *DSP1* function has been shown to cause homeotic transformations, similar to those observed in mutants of *Scr* and Hox genes of the Bithorax Complex, and it appears to regulate several pathways necessary for proper growth and development (Decoville et al., 2008; Mosrin-Huaman et al., 1998; Salvaing et al., 2006). Its expression in weevils appears to be complex, as different transcripts are shown to be up-regulated in both *S. oryzae* and *D. ponderosae*. It is possible that these expression differences are specifically associated with *Scr* and *dorsal* expression in the head.

### S. oryzae Scr-RNAi

Owing to the presence of a distinct rostrum in *S. oryzae* and a lack thereof in *D. ponderosae* (including difficulties in laboratory rearing of this species), RNAi experiments were conducted in the former, despite its relatively low expression of *Scr* compared to *D. ponderosae*. In comparing the *S. oryzae Scr*-RNAi phenotype (Fig. 4E-F) to that of the wild type (Fig. 4A-D), malformations are immediately visible in the head and prothorax. Upon closer inspection, homeotic transformations are seen in the labium and prothorax. In the labium, partial maxillary identity is acquired in which the prementum largely takes on what appears to be maxillary stipes morphology and the nearly absent palpus of the wild-type (reduced to only the small apical palpomere; Fig. 5J) is identically converted to the apical palpomere of the maxilla, including the lateral and apical sensilla (Fig. 5K). The prothorax partially adopts mesothoracic identity, in which the mesonotum is partially replicated and visible on the posterior part of the pronotum. The prothorax also forms a slight signature of the mesothoracic elytron, discernible as a projecting edge along the postero-lateral margins of the pronotum.

**Fig. 4.**
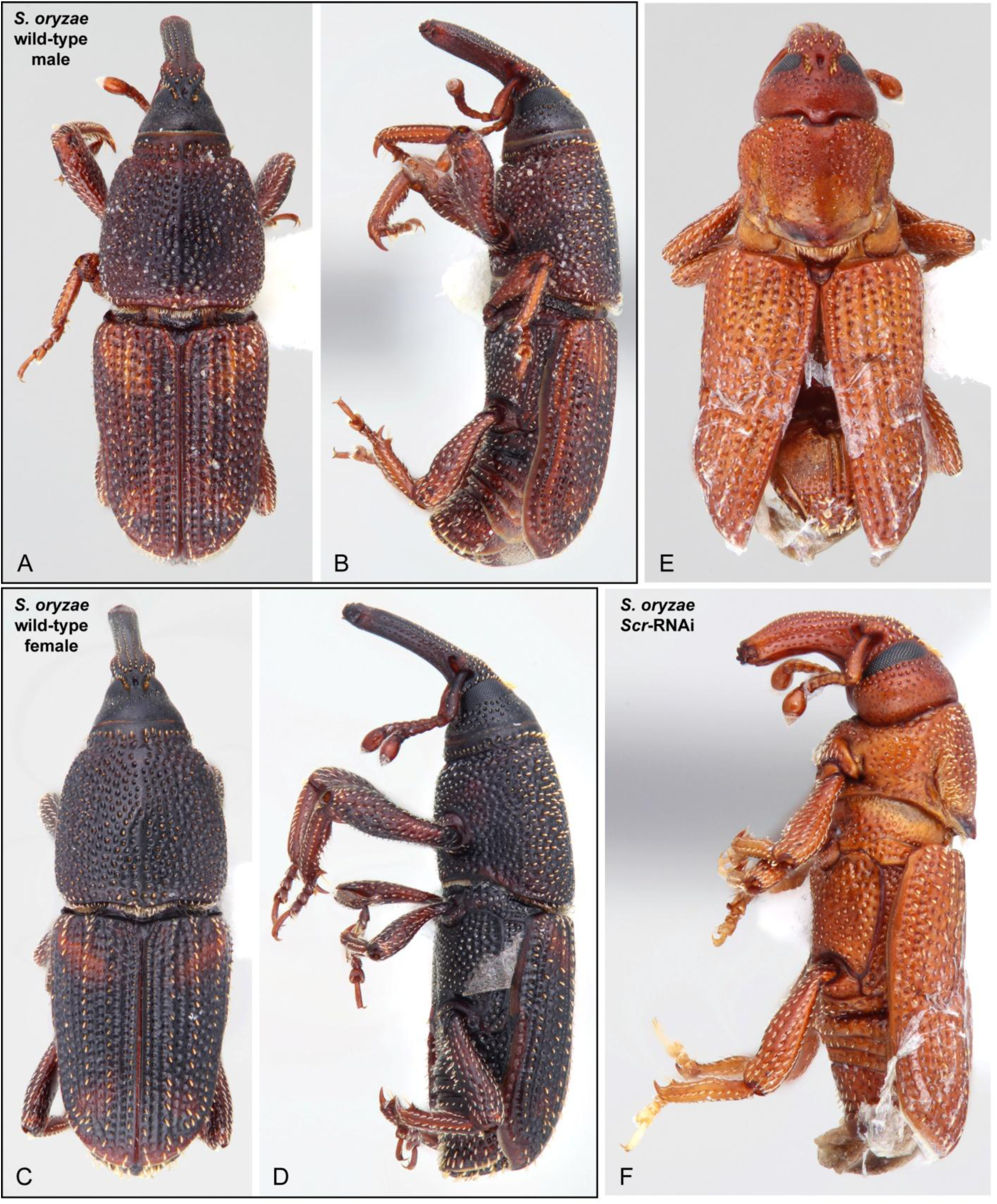
Wild-type of *Sitophilus oryzae*. **A-B**, male; **C-D**, female. *Scr*-RNAi phenotype of adult male. **E**, dorsal aspect; **F**, lateral aspect.

**Fig. 5.**
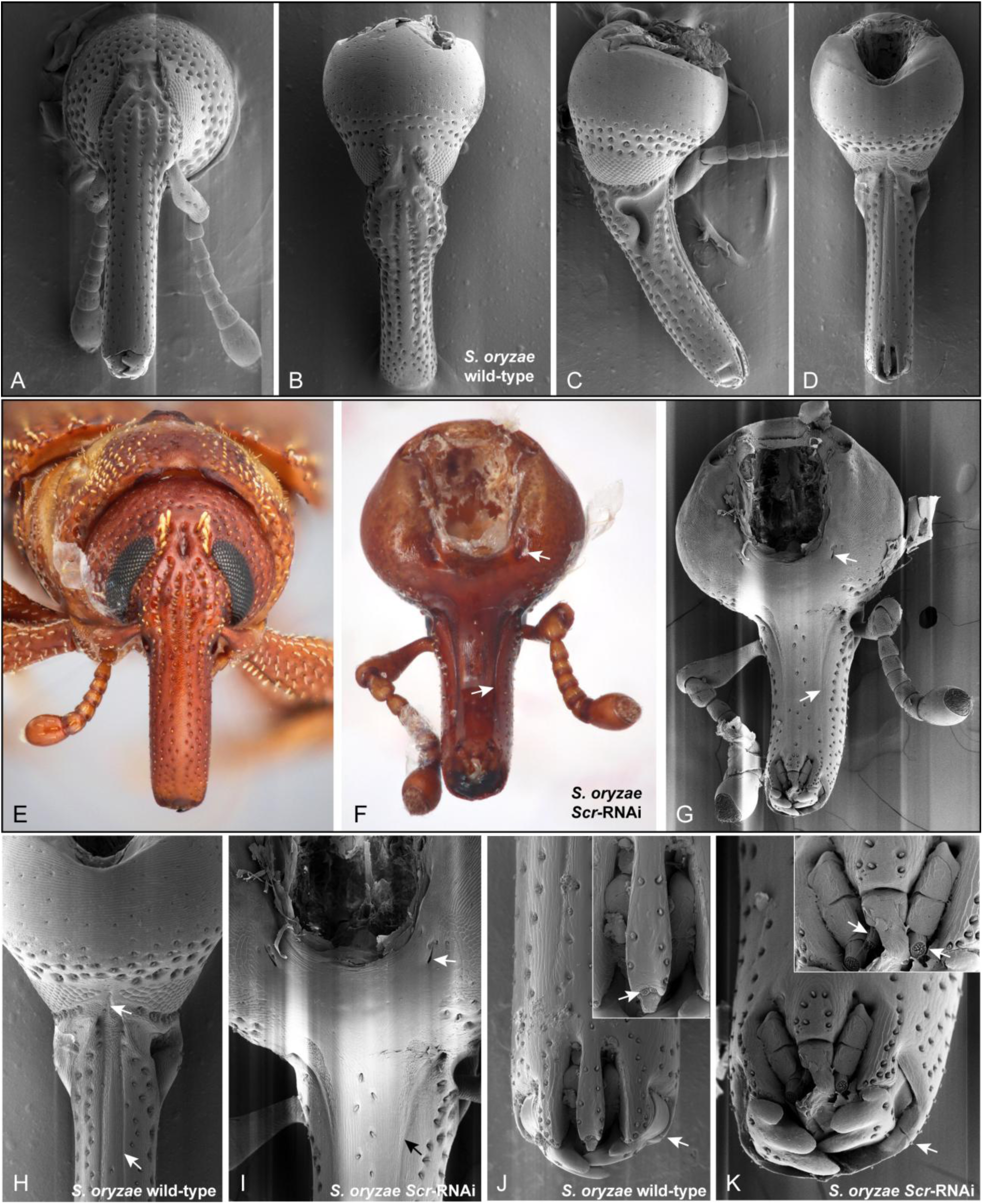
Wild-type adult head aspects of *S. oryzae* (**A-D, H, J**; SEM’s) and Scr-RNAi adult male head (**E-G, I, K**). **A**, anterior aspect of head; **B**, antero-dorsal aspect of head; **C**, lateral aspect of head; **D**, ventral aspect of head; **E**, anterior aspect of head; **F**, ventral aspect of head, arrows pointing to reformed posterior tentorial pit and subgenal sulcus; **G**, ventral aspect of head (SEM), arrows pointing to reformed posterior tentorial pits and subgenal sulcus; **H**, enlargement of head and basal region of rostrum in D, arrows pointing to single medial posterior tentorial pit and subgenal sulcus; **I**, enlargement of head and basal region of rostrum in G, arrows pointing to reformed posterior tentorial pits and subgenal sulcus; **J**, apical region of rostrum in D, arrow pointing to frontal sulcus; inset showing enlargement of labium and maxillae, arrow pointing to reduced labial palpi; **K**, apical region of rostrum in G, arrow pointing to frontal sulcus; inset showing enlargement of labium and maxillae, arrows pointing to apical maxillary palpomere and transformed labial palpus of close identity to apical maxillary palpomere.

The most readily apparent malformation occurs in the focal area in this study, the head and rostrum. Comparison of the wild-type heads (Fig. 5A-D, H, J) and those of *Scr*-RNAi individuals (Fig. 5E-G, I, K) reveals a shorter, stouter, and more curved rostrum of the latter. In anterior aspect (Fig. 5E), the RNAi rostrum is not significantly different, only slightly shorter and wider; however, the rostrum is markedly different in ventral aspect (Fig. 5F-G). The wild-type eyes are nearly confluent and touch ventrally, but they are not even visible ventrally in the *Scr*-RNAi head owing to an enlarged medial region along the entire ventral surface of the head. Anteriorly, the region that is probably the mentum is wider and shortened to approximately half of the wild-type length. Remnants of the frontal sulci are slightly more visible in the knockdown (Fig. 5K). Posteriorly, as partially delimited by distinctly separate posterior tentorial pits, is a large gular region that has reappeared just anterior to the occipital foramen. The two lateral pits are confirmed to be associated with the tentorium from histological sectioning and dissection of the head, showing separate arms of the posterior tentorium (Fig. 6F). In the wild-type, due to the loss of the gula, the arms of the posterior tentorium become fused ventrally; however, different from most weevils that have lost the gula and possess a single continuous gular suture extending from the occipital foramen to the posterior tentorial pits, much of the posterior tentorium is internalized in *Sitophilus*, thereby eliminating most of the suture and only connecting to the ventral head cuticle near the occipital foramen and medially between the eyes (Fig. 6C). In the heads of *Scr*-RNAi individuals, although the posterior tentorial connection between the eyes is lost, the gula still appears to extend anteriorly to this region according to the posterior extension of the subgenal sulci and position of the eyes in comparison to the wild-type. The occipital sulci also have become more widely separated and are proximally divergent, following the spacing of the antennal sockets. Medial to the occipital sulci is a pair of longitudinal lines that have reformed. Histological sectioning and position indicate that this pair of sulci (and associated internal apodemes) must be the subgenal sulci. This observation is confirmed by the tendons supporting the internal apodemes, namely those of the maxilla in the case of the re-formed subgenal sulci (Fig. 6D-E). This altered condition in *S. oryzae Scr*-RNAi most closely resembles that found in the weevil subfamily Rhynchitinae (Attelabidae), followed by the weevil families Nemonychidae and possibly Anthribidae, in which the maxillary tendons are supported by the apodemes of the subgenal sulci (Davis, 2017).

**Fig. 6.**
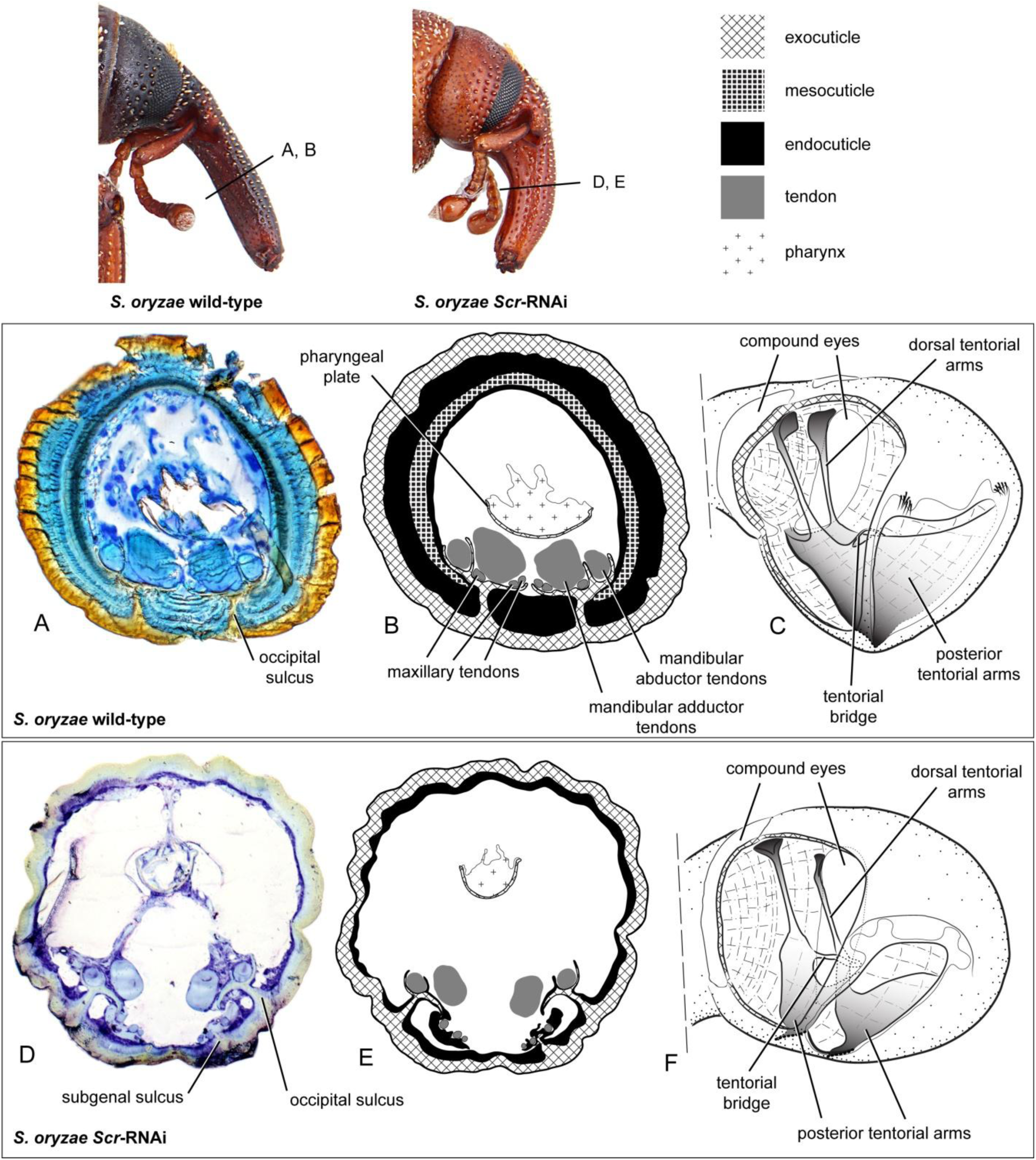
Semi-thin cross sections of *S. oryzae* wild-type (**A-B**) and *Scr*-RNAi (**D-E**) heads, and illustrations of wild-type (**C**) and *Scr*-RNAi (**F**) heads, highlighting tentorium. Top right displaying legend for diagrams. **A**, section at middle of rostrum, showing presence of one pair of apodemes emanating from occipital sulci; **B**, diagram of section in A; **C**, illustration of wild-type head, showing fused arms of posterior tentorium; **D**, section at middle of rostrum, showing presence of two pairs of apodemes emanating from both occipital and subgenal sulci; **E**, diagram of section in D; **F**, illustration of head of *Scr*-RNAi individual, showing presence of separated arms of the posterior tentorium.

**Fig. 7.**
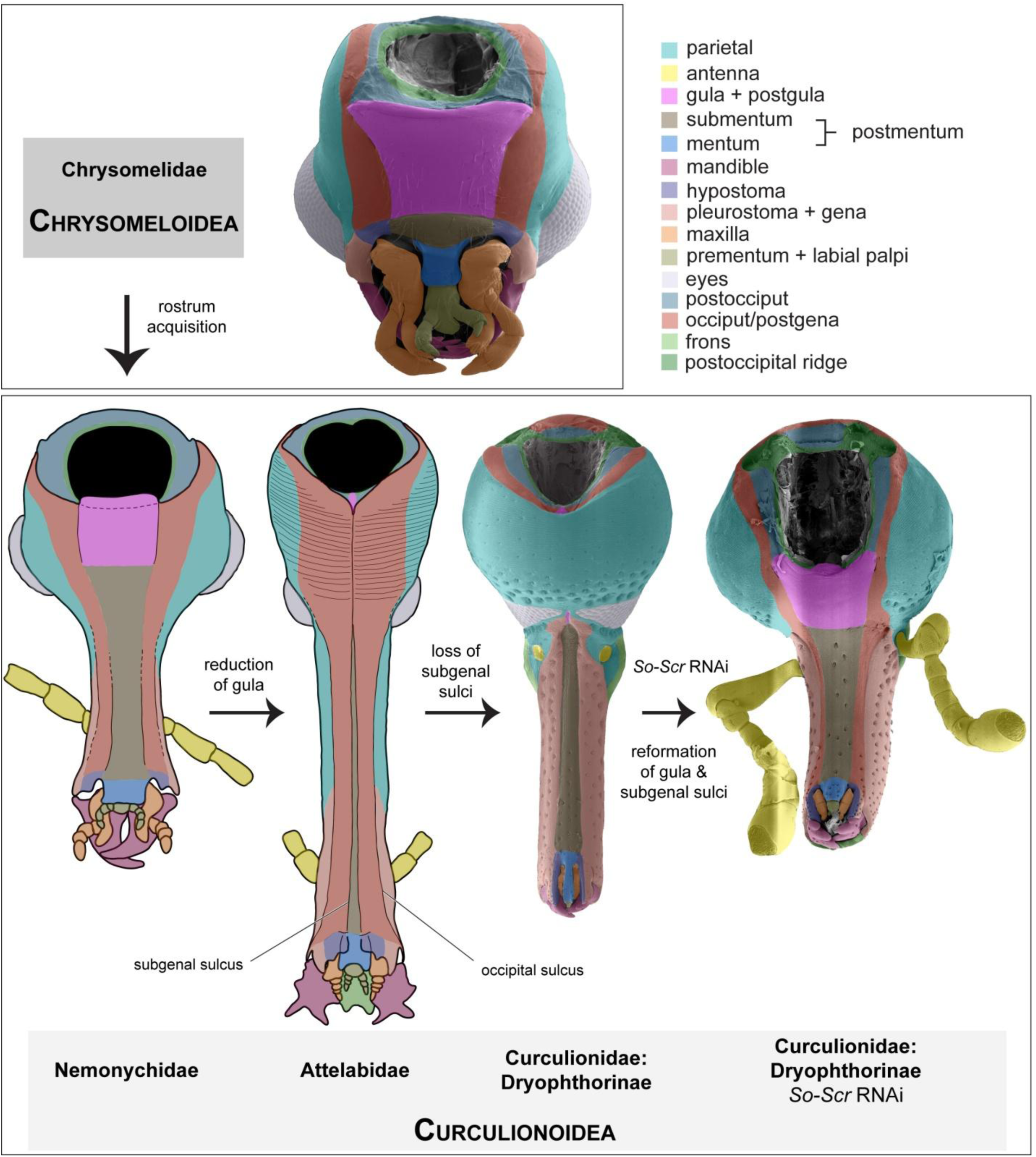
Head and rostrum structure and modification from an ancestral-like condition (as seen in Chrysomeloidea) to basal (Nemonychidae) and progressively more derived lineages (Attelabidae, Curculionidae) in Curculionoidea. Structural modification of the head and rostrum are also depicted in the head of *S. oryzae Scr*-RNAi individual. All heads are viewed in ventral aspects; heads of Nemonychidae and Attelabidae are illustrations, those of Chrysomelidae and *S. oryzae* are SEM’s. *So*=*Sitophilus oryzae*.

Excluding Nemonychidae, Anthribidae, and many Belidae, all remaining curculionoid lineages have lost the gula. Furthermore, all lineages, excepting Nemonychidae, Anthribidae, and Attelabidae have lost the subgenal sulci, although the lines sometimes are indiscernible externally in the former two families (Davis, 2017). In groups that have lost the subgenal sulci, the apodemes of the occipital sulci support the maxillary tendons in addition to those of the mandible (Fig. 6A-B). The fundamental trifurcate structure of the occipital apodemes was not modified in the *Scr*-RNAi head — the shape only slightly warped — suggesting that the subgenal apodemes indeed have been lost, as opposed to becoming fused with the occipital apodemes as in at least Dryophthorinae and perhaps all curculionoids that lack the subgenal sulci.

In addition to the above structural modifications, a few others are perhaps noteworthy as viewed from the semi-thin sections of the *Scr*-RNAi rostrum. First, the pharynx is smaller, including the pharyngeal plate supporting it that is composed of exocuticle (Fig. 6D-E), and it has shifted more dorsally within the rostrum. Second, the cuticle is thinner, including the exocuticle but particularly the endocuticle. The separation of these two cuticular layers is less distinct and any trace of mesocuticle also is absent. It is known that many (probably most) teneral adult holometabolous insects continue deposition of the endocuticle after eclosion, though the duration and extent of deposition varies. As the adult *S. oryzae Scr*-RNAi individuals did not survive long, it is possible that complete endocuticular deposition did not occur in the observed individuals. This latter scenario is a likely explanation given that the entire body also was relatively untanned, but it does not account for the indistinct separation of the endo- and exocuticles, as endocuticular deposition was well underway at this teneral stage (as evidenced by the thin endocuticular layer in the semi-thin sections) and a distinct separation of the two layers should have been evident.

### Scr function in Curculionoidea

The regulation and functions of *Scr* are still somewhat nebulous due to its complex interactions with and genomic proximity to *Antennapedia, fushi tarazu*, and *Deformed*. It is a Homeobox gene in the Antennapedia complex and is also well-known for its suppression of dorsal appendages on the prothorax (wing rudiments). In *S. oryzae Scr*-RNAi, this function in the prothorax is also visible. The resulting knockdown phenotype in the head and rostrum is interesting, in that the ventral surface of the head widens, caused by a re-formation of the gula, subgenal sulci, and intervening regions. At the rostral apex, the frontal sulci are slightly more formed. Aside from gular enlargement and subgenal sulcus re-formation, the modifications mentioned above are typical of *Scr* knockdown in *Tribolium, Drosophila*, and other insects (where the labium of the adult head is present; Curtis et al., 2001; Haas and Beeman, 2012; Hughes and Kaufman, 2000, 2002; Percival-Smith et al., 1997; Rogers et al., 1997, 2002; Smith and Jockusch, 2014). As for the other modifications, such changes likely do not occur in *Tribolium* because there have been few evolutionary alterations in regions posterior to the labium from the generalized beetle head groundplan.

Gular loss has occurred multiple times in Curculionoidea. As mentioned, in the basal weevil families Nemonychidae, Anthribidae, and many members of Belidae, the gula is retained but its delimitation is less pronounced. In Belidae, many belines retain a narrow gula but oxycorynines have lost it like the remaining curculionid families. Therefore, it is possible that trace expression of *Scr* in the gula and submentum began early in weevil evolution, some point at which it gained a repressive function in the formation of these regions, culminating at an expression threshold that eliminates parts of these regions entirely. Although gene expression was not examined *in situ* in this study, it is assumed that *Scr* protein expression is in close proximity to the anatomical regions that were affected by RNAi. This gain of repression of the postero-ventral head regions likely entails changes in *Scr* target proteins and mirrors its repressive function in other body regions, such as in its supression of dorsal appendages on the prothorax and protibial setal combs in some groups (Popadić et al., 1998). It is interesting to note that the relative expression level of *Scr* in *D. ponderosae* is markedly higher than in *S. oryzae*, yet the fundamental arrangement of the ventral head sulci is not drastically different. Aside from the absence of a rostrum, the gula in *D. ponderosae* also has been lost. Given the repression by *Scr* of fairly large ventral head structures in *S. oryzae*, it will be interesting to see the function of *Scr* in a scolytine (which exhibits a comparatively higher level of *Scr* expression) and if it demonstrates other unknown repressive functions related to the rostrum.

## Conclusion

Analyses of differential gene expression can be challenging among taxonomically disparate organisms due to wide evolutionary gaps between clades and corresponding divergent gene sequences. Nonetheless, a number of genes have been identified as promising candidates. This study illustrates the utility of transcriptomics across phylogenetic distances that span ∼110 mya in identifying genes responsible for complex adaptive structures. The preliminary comparisons between a rostrate weevil, *S. oryzae*, and a weevil that has evolutionarily lost its rostrum, *D. ponderosae*, illuminate some of the key genes involved in the formation and modification of the hallmark feature of weevils, the rostrum.

Future work will entail continuing RNAi experiments with interesting candidate genes and examining their spatial expression *in situ*. Gene expression and RNAi data would need to be compared to form hypotheses regarding the evolution of developmental pathways involved in rostrum formation. Examination of rostrum development, as with any morphological features, also will have remarkable influence in understanding homology statements when the relevant genetic elements involved in forming those structures are analyzed in comparison (Prud’homme et al., 2006).Together, these data will provide an extensive foundation from which to begin further research, including examination of these developmental genes in a broader phylogenetic context.

## Acknowledgements

Sincere thanks goes to Paulyn Cartwright (University of Kansas), as well as current and previous members of her lab, including Mariya Shcheglovitova, Annalise Nawrocki, Steve Sanders, and Sally Chang, for assisting in various aspects of Illumina sequencing and transcriptome analyses. I am grateful to the following people for their help in acquiring specimens of certain species: Jose Negron (USFS, Rocky Mountain Research Station) in acquiring *D. ponderosae* specimens of various developmental stages and James Throne (USDA, ARS, Grain Marketing & Production Research Center) in acquiring the starter colony for *S. oryzae*. Much appreciation goes to Susan Brown (Kansas State University) for her assistance in learning developmental and molecular techniques. Unending gratitude also goes to Michael S. Engel (Univ. of Kansas) for PhD dissertation guidance and to David Grimaldi (AMNH) for munificent postdoctoral support and greatly improving the manuscript. Cordial gratitude also goes to Gary Struhl (Columbia Univ.) for access to his laboratory during components of the study.

## Data availability

Raw sequence data and assembled transcriptomes for *Dendroctonus ponderosae* (BioSample) and *Sitophilus oryzae* (BioSample SAMN08668856) are publicly available through NCBI databases (both included under BioProject PRJNA437510). Raw sequence data is available through the Sequence Read Archive (SRA), including accessions SRR6830978 (*D. ponderosae*) and SRR6830122 (*S. oryzae*), and assembled transcriptomes are available through the Transcriptome Shotgun Assembly sequence database (TSA), including accessions XX (*D. ponderosae*) and XX (*S. oryzae*).

